# Replication-competent vesicular stomatitis virus vaccine vector protects against SARS-CoV-2-mediated pathogenesis

**DOI:** 10.1101/2020.07.09.196386

**Authors:** James Brett Case, Paul W. Rothlauf, Rita E. Chen, Natasha M. Kafai, Julie M. Fox, Swathi Shrihari, Broc T. McCune, Ian B. Harvey, Brittany Smith, Shamus P. Keeler, Louis-Marie Bloyet, Emma S. Winkler, Michael J. Holtzman, Daved H. Fremont, Sean P.J. Whelan, Michael S. Diamond

## Abstract

Severe acute respiratory syndrome coronavirus 2 (SARS-CoV-2) has caused millions of human infections and hundreds of thousands of deaths. Accordingly, an effective vaccine is of critical importance in mitigating coronavirus induced disease 2019 (COVID-19) and curtailing the pandemic. We developed a replication-competent vesicular stomatitis virus (VSV)-based vaccine by introducing a modified form of the SARS-CoV-2 spike gene in place of the native glycoprotein gene (VSV-eGFP-SARS-CoV-2). Immunization of mice with VSV-eGFP-SARS-CoV-2 elicits high titers of antibodies that neutralize SARS-CoV-2 infection and target the receptor binding domain that engages human angiotensin converting enzyme-2 (ACE2). Upon challenge with a human isolate of SARS-CoV-2, mice expressing human ACE2 and immunized with VSV-eGFP-SARS-CoV-2 show profoundly reduced viral infection and inflammation in the lung indicating protection against pneumonia. Finally, passive transfer of sera from VSV-eGFP-SARS-CoV-2-immunized animals protects naïve mice from SARS-CoV-2 challenge. These data support development of VSV-eGFP-SARS-CoV-2 as an attenuated, replication-competent vaccine against SARS-CoV-2.

## INTRODUCTION

Severe acute respiratory syndrome coronavirus 2 (SARS-CoV-2), a positive-sense, single-stranded, enveloped RNA virus, is the causative agent of coronavirus disease 2019 (COVID-19). Since its outbreak in Wuhan, China in December, 2019, SARS-CoV-2 has infected millions of individuals and caused hundreds of thousands of deaths worldwide. Because of its capacity for human-to-human transmission, including from asymptomatic individuals, SARS-CoV-2 has caused a pandemic, leading to significant political, economic, and social disruption (Bai et al., 2020). Currently, social quarantine, physical distancing, and vigilant hand hygiene are the only effective preventative measures against SARS-CoV-2 infections. Thus, effective countermeasures, particularly vaccines, are urgently needed to curtail the virus spread, limit morbidity and mortality, and end the COVID-19 pandemic.

The SARS-CoV-2 spike (S) protein mediates the receptor-binding and membrane fusion steps of viral entry. The S protein also is the primary target of neutralizing antibodies (Baum et al., 2020; Chi et al., 2020; Pinto et al., 2020; Rogers et al., 2020) and can elicit CD4^+^ and CD8^+^ T cell responses (Grifoni et al., 2020). Several SARS-CoV-2 vaccine platforms based on the S protein are being developed, including adenovirus-based vectors, inactivated virus formulations, recombinant subunit vaccines, and DNA- and mRNA-based strategies (Amanat and Krammer, 2020; Lurie et al., 2020). While several of these vaccines have entered human clinical trials, efficacy data in animals has been published for only a subset of these candidates (Gao et al., 2020; Yu et al., 2020).

We recently reported the generation and characterization of a replication-competent, VSV (designated VSV-eGFP-SARS-CoV-2) that expresses a modified form of the SARS-CoV-2 spike (Case et al., 2020). We demonstrated that monoclonal antibodies, human sera, and soluble ACE2-Fc potently inhibit VSV-eGFP-SARS-CoV-2 infection in a manner nearly identical to a clinical isolate of SARS-CoV-2. This suggests that chimeric VSV displays the S protein in an antigenic form that resembles native infectious SARS-CoV-2. Because of this data, we hypothesized that a replicating VSV-eGFP-SARS-CoV-2 might serve as an alternative platform for vaccine development. Indeed, an analogous replication-competent recombinant VSV vaccine expressing the Ebola virus (EBOV) glycoprotein protects against lethal EBOV challenge in several animal models (Garbutt et al., 2004; Jones et al., 2005), is safe in immunocompromised nonhuman primates (Geisbert et al., 2008), and was approved for clinical use in humans after successful clinical trials (Henao-Restrepo et al., 2017; Henao-Restrepo et al., 2015). Other live-attenuated recombinant VSV-based vaccines are in pre-clinical development for HIV-1, hantaviruses, filoviruses, arenaviruses, and influenza viruses (Brown et al., 2011; Furuyama et al., 2020; Garbutt et al., 2004; Geisbert et al., 2005; Jones et al., 2005).

Here, we determined the immunogenicity and *in vivo* efficacy of VSV-eGFP-SARS-CoV-2 as a vaccine candidate in a mouse model of SARS-CoV-2 pathogenesis. We demonstrate that a single dose of VSV-eGFP-SARS-CoV-2 generates a robust neutralizing antibody response that targets both the SARS-CoV-2 spike protein and the receptor binding domain (RBD) subunit. Upon challenge with infectious SARS-CoV-2, mice immunized with one or two doses of VSV-eGFP-SARS-CoV-2 showed significant decreases in lung and peripheral organ viral loads, pro-inflammatory cytokine responses, and consequent lung disease. VSV-eGFP-SARS-CoV-2-mediated protection likely is due in part to antibodies, as passive transfer of immune sera to naïve mice limits infection after SARS-CoV-2 challenge. This study paves the way for further development of a VSV-vectored SARS CoV-2 vaccine.

## RESULTS

### eneration of a VSV-eGFP-SARS-CoV-2 as a vaccine platform

We previously reported a chimeric, replication-competent VSV expressing the SARS-CoV-2 spike protein as an effective platform for measuring neutralizing antibodies (Case et al., 2020). As replication-competent VSVs are in clinical use as vaccines for emerging RNA viruses or in pre-clinical development (Fathi et al., 2019), we tested whether VSV-eGFP-SARS-CoV-2 could protect mice against SARS-CoV-2.

To examine the immune response to VSV-eGFP-SARS-CoV-2, we immunized four-week-old BALB/c mice with 10^6^ plaque-forming units (PFU) of VSV-eGFP-SARS-CoV-2 or a control, VSV-eGFP (**Fig 1A**). As murine ACE2 does not serve as a receptor for SARS-CoV-2, we spiked our preparation of VSV-eGFP-SARS-CoV-2 with trace amounts of VSV G to permit a single round of infection, an approach used previously for SARS-CoV (Kapadia et al., 2008). At 28 days post-priming, one cohort of animals was boosted with the homologous vaccine. Serum was isolated from all animals at three weeks post priming or boosting, and IgG titers against recombinant SARS-CoV-2 S protein or the RBD were determined by ELISA (**Fig 1B-C**). Immunization with VSV-eGFP-SARS-CoV-2 induced high levels of anti-S and anti-RBD-specific IgG compared to control VSV-eGFP with reciprocal median serum endpoint titers of 3.2 × 10^5^ and 2.7 × 10^6^ (anti-S) and 1.1 × 10^4^ and 1.4 × 10^4^ (anti-RBD) for one and two doses of vaccine, respectively.

**Figure 1.**
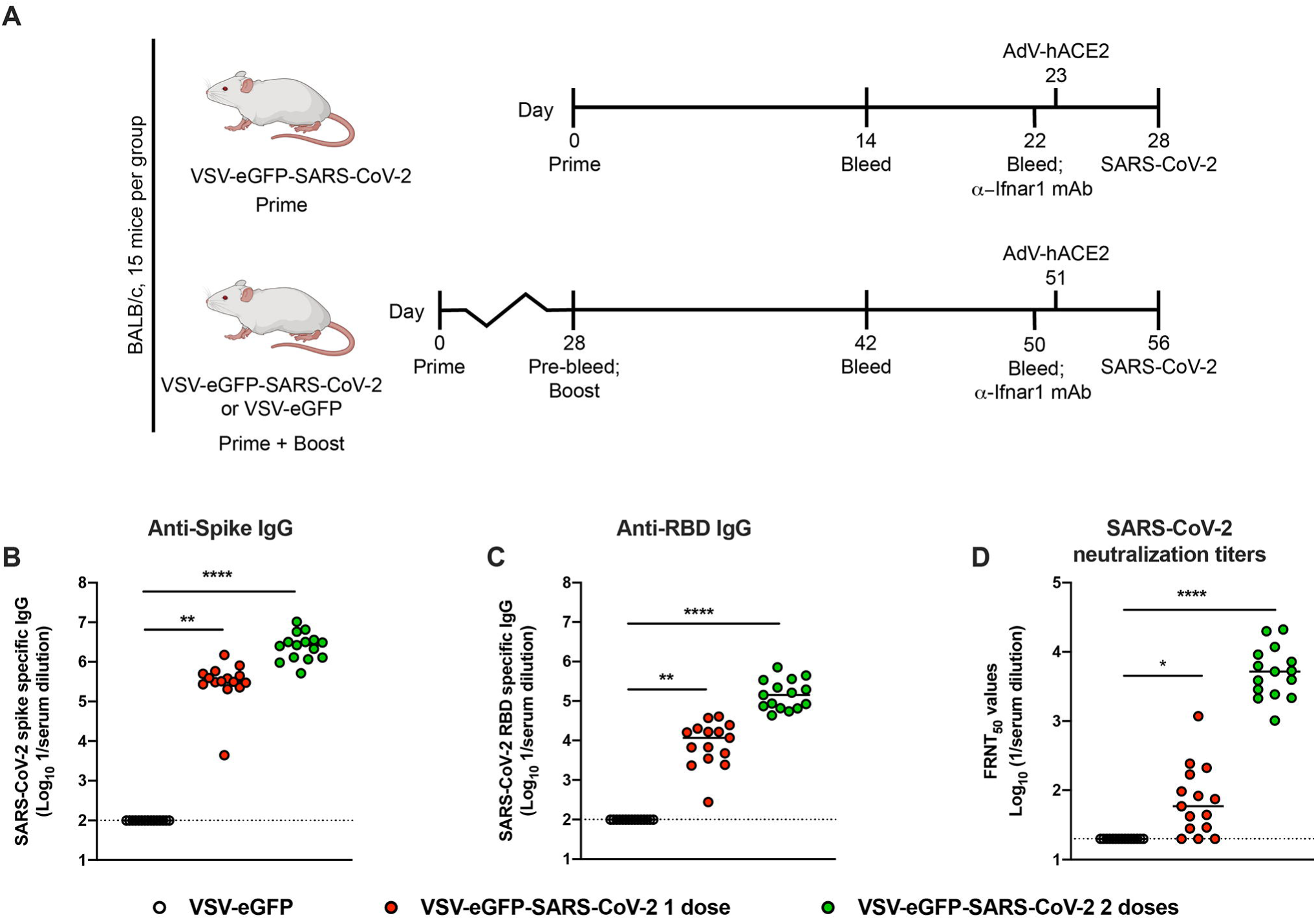
Immunogenicity of VSV-eGFP-SARS-CoV-2. **A**. Scheme of vaccination and SARS-CoV-2 challenge. **B-D**. Four-week-old female BALB/c mice were immunized with VSV-eGFP or VSV-eGFP-SARS-CoV-2. Some of the immunized mice were boosted with their respective vaccines four weeks after primary vaccination. IgG responses in the sera of vaccinated mice were evaluated three weeks after priming or boosting by ELISA for binding to SARS-CoV-2 S (**B**) or RBD (**C**) or two weeks after priming or boosting by focus reduction neutralization test (FRNT) (**D**) (n = 15 per group; one-way ANOVA with Dunnett’s post-test: **** *P* < 0.0001).

We measured neutralizing antibody titers against SARS-CoV-2 after priming or boosting using a focus-reduction neutralization test (Case et al., 2020). Immunization with a single or two-dose regimen of VSV-eGFP-SARS-CoV-2 induced neutralizing antibodies (median titers of 1/59 and 1/5206, respectively) whereas the control VSV-eGFP vaccine did not (**Fig 1D**). Boosting was effective and resulted in a 90-fold increase in neutralizing activity after the second dose of VSV-eGFP-SARS-CoV-2. Collectively, these data suggest that VSV-eGFP-SARS-CoV-2 is immunogenic and elicits high titers of antibodies that neutralize infection and target the RBD of the SARS-CoV-2 S protein.

### VSV-eGFP-SARS-CoV-2 protects mice against SARS-CoV-2 infection

Four weeks after priming or priming and boosting, mice were challenged with SARS-CoV-2 (strain 2019 n-CoV/USA_WA1/2020) after delivery of a replication-defective adenovirus expressing human ACE2 (AdV-hACE2) that enables receptor expression in the lungs (Hassan et al., 2020). Immunized mice were administered 2 mg of anti-Ifnar1 mAb one day prior to intranasal delivery of AdV-hACE2. We administer anti-Ifnar1 antibody to augment virus infection and create a stringent disease model for vaccine protection. Five days later, mice were inoculated with 3 × 10^5^ PFU of SARS-CoV-2 via the intranasal route (**Fig 1A**) and subsequently, we measured viral yield by plaque and RT-qPCR assays. At day 4 post-infection (dpi) infectious virus was not recovered from lungs of mice vaccinated either with one or two doses of VSV-eGFP-SARS-CoV-2 (**Fig 2A**). For mice receiving only one dose of VSV-eGFP-SARS-CoV-2 vaccine, we observed a trend towards decreased levels of viral RNA in the lung, spleen, and heart at 4 dpi and in the lung and spleen at 8 dpi compared to the control VSV-eGFP vaccinated mice (**Fig 2B-E**). Mice that received two doses of VSV-eGFP-SARS-CoV-2 had significantly lower levels of viral RNA in most tissues examined compared to control VSV-eGFP vaccinated mice (**Fig 2B-E**). Consistent with our viral RNA measurements, we observed less SARS-CoV-2 RNA by *in situ* hybridization in lung tissues of VSV-eGFP-SARS-CoV-2 immunized mice at 4 dpi (**Fig 2F**). Collectively, these data support that immunization with VSV-eGFP-SARS-CoV-2 protects against SARS-CoV-2 infection in mice.

**Figure 2.**
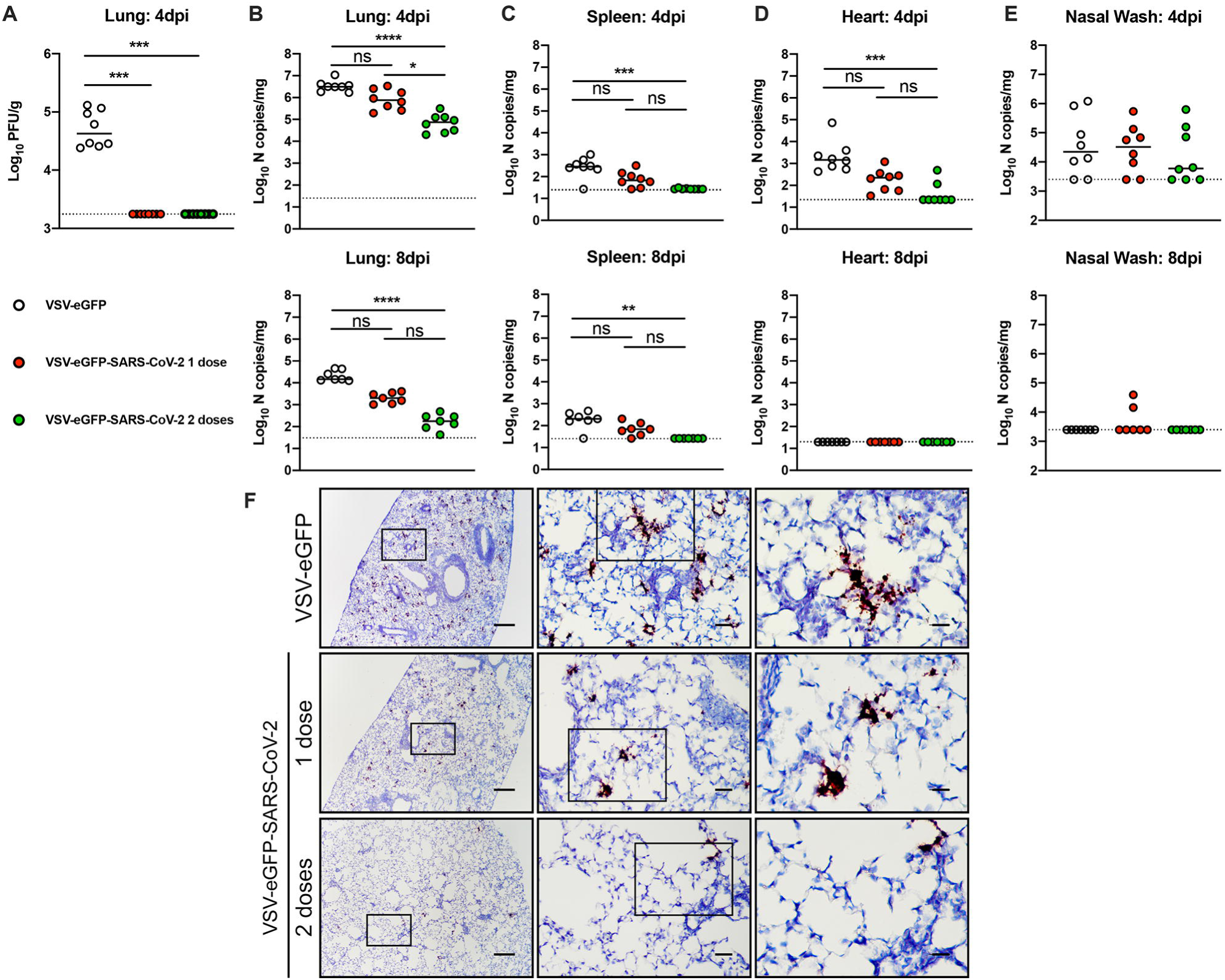
VSV-eGFP-SARS-CoV-2 protects mice against SARS-CoV-2 infection. Three weeks after priming or boosting with VSV-eGFP or VSV-eGFP-SARS-CoV-2, immunized animals were treated with anti-Ifnar1 mAb and one day later, animals were transduced with 2.5 × 10^8^ PFU of AdV-hACE2 by intranasal administration. Five days later, animals were challenged with 3 × 10^5^ PFU of SARS-CoV-2 via intranasal administration. **A-E**. At 4 or 8 dpi tissues were harvested and viral burden was determined in the lung (**A-B**), spleen (**C**), heart (**D**), and nasal washes (**E**) by plaque (**A**) or RT-qPCR (**B**-**E**) assay (n = 7-8 mice per group; Kruskal-Wallis test with Dunn’s post-test (**A-E**): ns, not significant, * *P* < 0.05, ** *P* < 0.01, *** *P* < 0.001, **** *P* < 0.0001). Dotted lines indicate the limit of detection. **F**. SARS-CoV-2 RNA *in situ* hybridization of lungs of mice vaccinated with VSV-eGFP or VSV-eGFP-SARS-CoV-2 and challenged with SARS-CoV-2 at 4 dpi. Images show low- (left; scale bars, 100 μm), medium- (middle; scale bars, 100 μm), and high-power magnifications (right; scale bars, 10 μm; representative images from n = 3 per group).

### VSV-eGFP-SARS-CoV-2 limits SARS-CoV-2-induced lung inflammation

Both SARS-CoV and SARS-CoV-2 typically cause severe lung infection and injury that is associated with high levels of pro-inflammatory cytokines and immune cell infiltrates (Gu and Korteweg, 2007; Huang et al., 2020). The AdV-hACE2 transduced mouse model of SARS-CoV-2 pathogenesis recapitulates several aspects of lung inflammation and coronavirus disease (Hassan et al., 2020). To assess whether VSV-eGFP-SARS-CoV-2 limits virus-induced inflammation, we measured pro-inflammatory cytokine and chemokine mRNA in lung homogenates from vaccinated animals at 4 dpi by RT-qPCR assays (**Fig 3A**). Animals immunized with one or two doses of VSV-eGFP-SARS-CoV-2 had significantly lower levels of pro-inflammatory cytokine and chemokine mRNA compared to VSV-eGFP vaccinated mice. Specifically, type I and III interferons (IFN-β and IFN-λ) were decreased early during infection in both one-dose and two-dose groups of mice immunized with VSV-eGFP-SARS-CoV-2. While there were no detectable differences in IFN-γ or TNF-α levels between groups, IL-6 and IL-1β were lower at 4 dpi after VSV-eGFP-SARS-CoV-2 vaccination. Similarly, levels of mRNAs encoding chemokines CXCL1, CXCL10, and CXCL11, which recruit immune cells to the lung, were decreased at 4 dpi in VSV-eGFP-SARS-CoV-2 compared to VSV-eGFP immunized mice.

**Figure 3.**
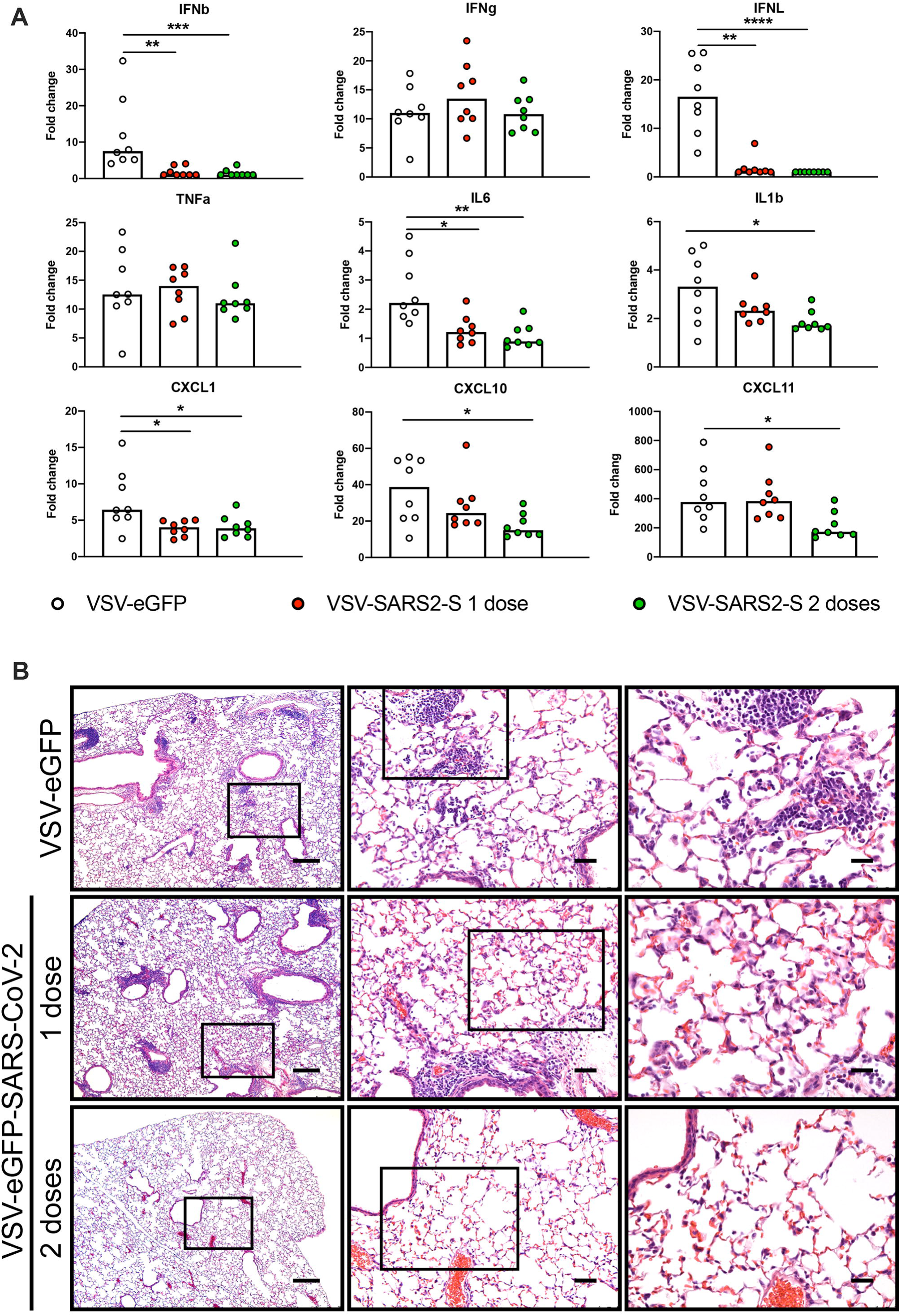
VSV-eGFP-SARS-CoV-2 protects mice from SARS-CoV-2 lung inflammation A. Lungs of VSV-eGFP or VSV-eGFP-SARS-CoV-2 immunized mice were evaluated at 4 dpi for cytokine and chemokine expression by RT-qPCR assay. Data are shown as fold-change in gene expression compared to fully naïve, age-matched animals after normalization to *Gapdh* (n = 7-8 per group, Kruskal-Wallis test with Dunn’s post-test: * *P* < 0.05, ** *P* < 0.01, *** *P* < 0.001, **** *P* < 0.0001). **B**. Hematoxylin and eosin staining of lung sections from immunized mice at 8 dpi with SARS-CoV-2 (3 × 10^5^ PFU). Images show low- (left; scale bars, 250 μm), medium- (middle; scale bars, 50 μm), and high-power magnifications (right; scale bars, 25 μm; representative images from n = 3 mice per group).

To determine the extent of lung pathology in SARS-CoV-2 challenged mice, at 8 dpi, we stained lung sections with hematoxylin and eosin (**Fig 3B**). Lung sections from VSV-eGFP-immunized mice showed immune cell (including neutrophil) infiltration into perivascular, peribronchial, and alveolar locations consistent with viral pneumonia. Lung sections from mice immunized with one dose of VSV-eGFP-SARS-CoV-2 also showed some signs of inflammation. However, mice immunized with two doses of VSV-eGFP-SARS-CoV-2 showed substantially less accumulation of inflammatory cells at the same time point after SARS-CoV-2 infection. These data suggest that immunization with VSV-eGFP-SARS-CoV-2 generates a protective immune response, which limits SARS-CoV-2-induced lung disease in mice. In this model, two sequential immunizations show greater efficacy than a single one.

### Vaccine-induced sera limits SARS-CoV-2 infection

To investigate the contribution of antibodies in vaccine-mediated protection, we performed passive transfer studies. Serum was collected from VSV-eGFP and VSV-eGFP-SARS-CoV-2 vaccinated mice after one or two immunizations. Ten-week-old female BALB/c mice were administered anti-Ifnar1 mAb and AdV-hACE2 as described above to render animals susceptible to SARS-CoV-2. Five days later, 100 μL of pooled immune or control sera was administered by intraperitoneal injection. One day later, mice were inoculated with 3 × 10^5^ PFU of SARS-CoV-2 via the intranasal route (**Fig 4A**). Passive transfer of sera from animals vaccinated with VSV-eGFP-SARS-CoV-2 protected against SARS-CoV-2 infection compared to sera from the VSV-eGFP-immunized mice. At 4 dpi, lungs from animals treated with VSV-eGFP-SARS-CoV-2 immune sera from prime-only and boosted animals showed substantially reduced infectious virus burden (**Fig 4B**). Although not as striking, significant decreases in viral RNA levels also were observed in the lung and spleen of animals receiving VSV-eGFP-SARS-CoV-2 boosted sera compared to the VSV-eGFP sera (**Fig 4C-D**). Possibly, some of the viral RNA in lung tissue homogenates after passive transfer may represent neutralized virus within cells that has not yet been cleared. Viral RNA levels in the heart of animals given sera from VSV-eGFP-SARS-CoV-2 boosted mice trended toward, but did not reach, statistical significance (**Fig 4E**). No effect was observed in the nasal washes of any treated group (**Fig 4F**), consistent with the results from our vaccinated and challenged animals (**Fig 2E**).

**Figure 4.**
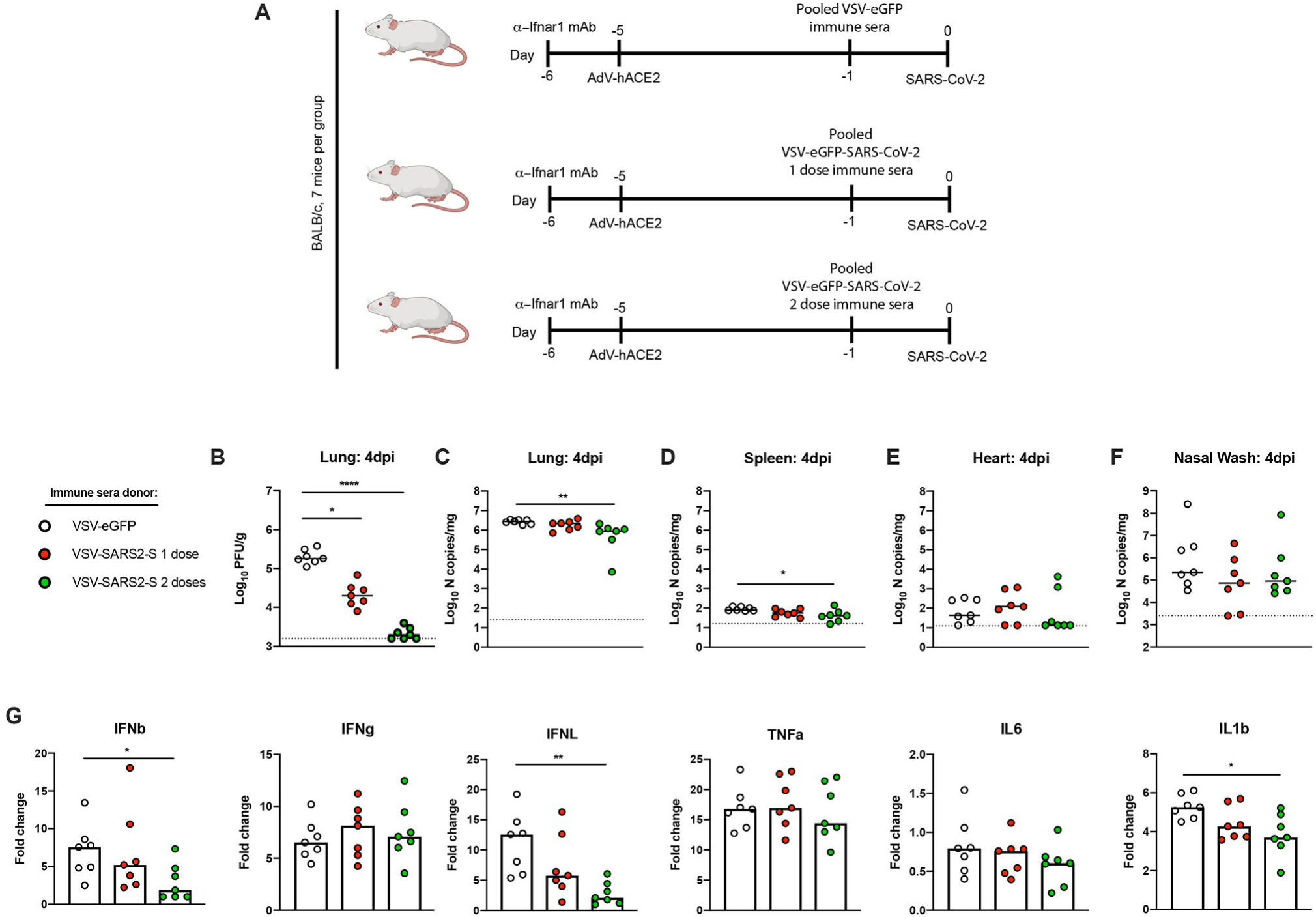
Vaccine-induced sera limits SARS-CoV-2 infection. **A**. Passive transfer of immune sera and SARS-CoV-2 challenge scheme. Ten-week-old female BALB/c mice were treated with anti-Ifnar1 mAb and one day later, animals were transduced with 2.5 × 10^8^ PFU of AdV-hACE2 by intranasal administration. Four days later, animals were administered 100 μL of pooled immune sera collected from VSV-eGFP or VSV-eGFP-SARS-CoV-2 vaccinated mice after one or two immunizations. One day later, animals were challenged with 3 × 10^5^ PFU of SARS-CoV-2 via intranasal administration. **B**-**F**. At 4 dpi tissues were harvested and viral burden was determined in the lung (**B**-**C**), spleen (**D**), heart (**E**), and nasal washes (**F**) by plaque (**B**) or RT-qPCR (**C**-**F**) assays (n = 7 mice per group; Kruskal-Wallis test with Dunn’s post-test (**B-F**): * *P* < 0.05, ** *P* < 0.01, **** *P* < 0.0001). Dotted lines indicate the limit of detection. **G**. Lungs of mice treated with immune sera were evaluated at 4 dpi for cytokine expression by RT-qPCR assay. Data are shown as fold-change in gene expression compared to naïve, age-matched animals after normalization to *Gapdh* (n = 7 per group, Kruskal-Wallis test with Dunn’s post-test: * *P* < 0.05, ** *P* < 0.01).

To determine the effect of the passive transfer of sera on SARS-CoV-2-mediated inflammation, we assessed the induction of several cytokines in the lung at 4 dpi (**Fig 4G**). Treatment with sera from animals immunized with two doses of VSV-eGFP-SARS-CoV-2 limited induction of some (IFN-β, IFN-λ, and IL-1β) pro-inflammatory cytokines after SARS-CoV-2 challenge. Together, these data suggest that antibodies are a major correlate of VSV-eGFP-SARS-CoV-2-mediated protection against SARS-CoV-2.

## DISCUSSION

The emergence of SARS-CoV-2 into the human population has caused a global pandemic, resulting in millions of infected individuals and hundreds of thousands of deaths. Despite initial indications that the pandemic had peaked, reopening of countries and renewed human-to-human contact has resulted in a recent surge in case numbers, suggesting that SARS-CoV-2 vaccines will be critical for curtailing the pandemic and resuming normal social interactions. In this study, we tested the efficacy of a replication-competent VSV-eGFP-SARS-CoV-2 vaccine. A single dose of VSV-eGFP-SARS-CoV-2 was sufficient to induce antibodies that neutralize SARS-CoV-2 infection and target the RBD and S protein, and a second dose substantially boosted this response. We then challenged mice with SARS-CoV-2 via the intranasal route and observed a complete loss of recovery of infectious virus in the lung in animals immunized with either one or two doses of VSV-eGFP-SARS-CoV-2. Compared to a single dose, administration of two doses of VSV-eGFP-SARS-CoV-2 elicited greater protection with further diminished viral loads. Immunization with VSV-eGFP-SARS-CoV-2 decreased the induction of several key pro-inflammatory cytokines and protected mice from alveolar inflammation, lung consolidation, and viral pneumonia. We also established an important role for protective antibodies, as passive transfer of immune sera from VSV-eGFP-SARS-CoV-2 immunized animals decreased viral burden and inflammation in the lung.

Recombinant VSV-based vaccines that encode viral glycoproteins have several advantages as a platform. Whereas DNA plasmid and mRNA-based vaccines have not yet been approved in the United States or elsewhere, Merck’s ERVEBO®, a replication-competent VSV expressing the EBOV glycoprotein, is currently in use in humans (Huttner et al., 2015). As a replicating RNA virus, VSV-based vaccines often can be used as single-dose administration and effectively stimulate both humoral and cellular immunity. Recombinant VSV grows efficiently in mammalian cell culture, enabling simple, large-scale production. Advantages of VSV as a vaccine vector also include the lack of homologous recombination and its non-segmented genome structure, which precludes genetic reassortment and enhances its safety profile (Lichty et al., 2004; Roberts et al., 1999). Unlike other virus-based vaccine vectors (Barouch et al., 2004; Casimiro et al., 2003; Santra et al., 2007), there is little preexisting human immunity to VSV as human infections are rare (Roberts et al., 1999) with the exception of some regions of Central America (Cline, 1976) or a limited number of at-risk laboratory workers (Johnson et al., 1966).

Several vaccine candidates for SARS-CoV-2 have been tested for immunogenicity. Our VSV-eGFP-SARS-CoV-2 vaccine elicited high levels of inhibitory antibodies with median and mean serum neutralizing titers of greater than 1/5,000. Two doses of VSV-eGFP-SARS-CoV-2 induced higher neutralizing titers with more rapid onset than similar dosing of an inactivated SARS-CoV-2 vaccine in the same strain of mice (Gao et al., 2020). Consistent with these results, serum anti-S endpoint titers were higher from mice immunized with two doses of VSV-eGFP-SARS-CoV-2 (1/2,700,000) than the highest two-dose regimen of the inactivated virion vaccine (1/820,000). Two doses of DNA plasmid vaccines encoding variants of the SARS-CoV-2 S protein induced relatively modest neutralizing antibody responses (serum titer of 1/170) in rhesus macaques. Related to this, anti-S titers were approximately 1,000-fold lower after two doses of the optimal DNA vaccine (Yu et al., 2020) when compared to two doses of VSV-eGFP-SARS-CoV-2. In a pre-print study, a single-dose of a chimpanzee adenovirus vectored vaccine encoding SARS-CoV-2 S protein, ChAdOx1 nCoV-19, also produced relatively low levels of serum neutralizing antibodies in mice and NHPs (1/40 to 1/80 in BALB/c and CD1 mice and < 1/20 in rhesus macaques). This data corresponded with anti-S1 and anti-S2 mean serum titers of between 1/100 and 1/1,000 in BALB/c mice and anti-S titers of < 1/1,000 in NHPs (DOI: 10.1101/2020.05.13.093195). Two doses of a recombinant adenovirus type-5 vectored SARS-CoV-2 vaccine in humans also produced relatively low RBD-binding (1/1,445 at day 28 post-boost) and neutralizing antibody (1/34 at day 28 post-boost) (Zhu et al., 2020). Finally, based on pre-print data (DOI: 10.1101/2020.06.11.145920), BALB/c mice immunized with two 1 µg doses of an mRNA vaccine candidate, mRNA-1273, elicited serum anti-S endpoint titers of 1/100,000. These mice produced mean neutralizing antibodies titers of approximately 1/1,000 and did not show evidence of infectious virus in the lung or nares after SARS-CoV-2 challenge.

Even though VSV-eGFP-SARS-CoV-2 is replication-competent and capable of spread, it likely did not do so efficiently in our BALB/c mice because the SARS-CoV-2 spike cannot efficiently utilize murine ACE2 for viral entry (Letko et al., 2020). This likely explains our need for boosting, as the response we observed likely was enabled by the residual small amount of *trans*-complementing VSV G to pseudotype the virions expressing the S protein in a manner similar to VSV-SARS (Kapadia et al., 2008), which effectively limited vaccine virus replication to a single cycle. We anticipate that in animals expressing ACE2 receptors competent for S binding, a single dose of VSV-eGFP-SARS-CoV-2 will be associated with greater immunogenicity, and not require a second immunization for protection. To test this hypothesis, immunization and challenge studies are planned in hACE2 transgenic mice (Bao et al., 2020; Jiang et al., 2020; Sun et al., 2020) as they become widely available, and in hamsters and NHPs.

Vaccine safety is a key requirement of any platform. Pathogenicity and immunogenicity of VSV is associated with its native glycoprotein G, which, in turn, determines its pan-tropism (Martinez et al., 2003). Replacing the glycoprotein of VSV with a foreign glycoprotein often results in virus attenuation *in vivo*. Indeed, the vast majority of cases where VSV recombinants express a heterologous viral glycoprotein (*e*.*g*., chikungunya virus, H5N1 influenza virus, Lassa virus, lymphocytic choriomeningitis virus, or Ebola virus) and were injected via intracranial route into mice or NHPs, no disease was observed (Mire et al., 2012; Muik et al., 2014; van den Pol et al., 2017; Wollmann et al., 2015). One exception is when VSV expressing the glycoproteins of the highly neurotropic Nipah virus was injected via an intracranial route into adult mice (van den Pol et al., 2017). Should substantial reactogenicity or neuronal infection be observed with VSV-eGFP-SARS-CoV-2, the vaccine could be attenuated further by introducing mutations into the matrix protein (Rabinowitz et al., 1981) or methyltransferase (Li et al., 2006; Ma et al., 2014), rearranging the order of genes (Ball et al., 1999; Wertz et al., 1998), or recoding of the L gene (Wang et al., 2015). The presence of the additional eGFP gene inserted between the leader and N genes also attenuates virus replication in cell culture (Whelan et al., 2000). Further development of a VSV vectored vaccine for SARS-CoV-2 likely will require deletion of eGFP from the genome, which may necessitate additional strategies of attenuation.

Future studies are planned to evaluate the durability of VSV-eGFP-SARS-CoV-2 and a variant lacking eGFP in inducing immunity. Other replication-competent VSV-based vaccines such as the rVSVΔG-ZEBOV-GP have been shown to generate long-lasting immune responses and protection (Kennedy et al., 2017). In addition, we plan to investigate in greater detail the contributions of additional arms of immunity in mediating protection. The robust induction of neutralizing antibodies elicited by one and two doses of VSV-eGFP-SARS-CoV-2 was a correlate of protection, as passive transfer of immune sera reduced viral infection and inflammation in the lung upon SARS-CoV-2 challenge. Nonetheless, it will be important to determine if additional immune responses, particularly CD8^+^ T cells, have an important protective role. Recently, SARS-CoV-2 specific CD4^+^ and CD8^+^ T cells were shown to be present in 100% and 70% of COVID-19 convalescent patients, respectively, with many of the T cells recognizing peptides derived from the S protein (Grifoni et al., 2020). Moreover, additional experiments are planned in aged animals (hACE2-expressing mice, hamsters, and NHPs) to address immunogenicity and protection in this key target population at greater risk for severe COVID-19. Overall, our data show that VSV-eGFP-SARS-CoV-2 can protect against severe SARS-CoV-2 infection and lung disease, supporting its further development as a vaccine.

## ACKNOWLEDGEMENTS

This study was supported by NIH contracts and grants (75N93019C00062, HHSN272201700060C and R01 AI127828, R37 AI059371 and U01 AI151810), the Defense Advanced Research Project Agency (HR001117S0019), R01 AI130591 and R35 HL145242, and gifts to Washington University. J.B.C. is supported by a Helen Hay Whitney Foundation postdoctoral fellowship. We thank Natalie Thornburg for providing the clinical isolate of SARS-CoV-2, Ahmed Hassan for amplifying the AdV-hACE2 stocks, and James Earnest for providing cell culture support. Some of the figures were created using BioRender.com.

## AUTHOR CONTRIBUTIONS

J.B.C. designed experiments, propagated the SARS-CoV-2 stocks, performed VSV immunizations, and SARS-CoV-2 challenge experiments. P.W.R. generated the VSV vaccines. J.B.C., R.E.C., N.M.K., J.M.F., S.S., and E.S.W. performed tissue harvests, histopathological studies, and viral burden analyses. B.T.M. performed in situ hybridization. I.B.H. and B.S. performed ELISAs. S.P.K. and M.J.H. analyzed the tissue sections for histopathology. J.B.C. and R.E.C. performed neutralization assays. L.M.B. generated the VSV-eGFP control. J.B.C., P.W.R., S.P.J.W., and M.S.D. wrote the initial draft, with the other authors providing editing comments.

## STAR METHODS

### RESOURCE AVAILABLITY

#### Lead Contact

Further information and requests for resources and reagents should be directed to and will be fulfilled by the Lead Contact, Michael S. Diamond.

#### Materials Availability

All requests for resources and reagents should be directed to and will be fulfilled by the Lead Contact author. This includes mice, antibodies, viruses, and proteins. All reagents will be made available on request after completion of a Materials Transfer Agreement.

#### Data and code availability

All data supporting the findings of this study are available within the paper and are available from the corresponding author upon request.

### EXPERIMENTAL MODEL AND SUBJECT DETAILS

#### Cells

BSRT7/5, Vero CCL81, Vero E6, Vero E6-TMPRSS2 (Case et al., 2020), and Vero-furin (Mukherjee et al., 2016) cells were maintained in humidified incubators at 34 or 37°C and 5% CO_2_ in DMEM (Corning) supplemented with glucose, L-glutamine, sodium pyruvate, and 10% fetal bovine serum (FBS).

#### Plasmids

The S gene of SARS-CoV-2 isolate Wuhan-Hu-1 (GenBank MN908947.3) was cloned into the backbone of the infectious molecular clone of VSV containing eGFP (pVSV-eGFP) as described (Case et al., 2020). pVSV-eGFP was used as previously described, but contains a mutation K535R, the phenotype of which will be described elsewhere. Expression plasmids of VSV N, P, L, and G were previously described (Stanifer et al., 2011; Whelan et al., 1995).

#### Recombinant VSV

VSV-eGFP-SARS-CoV-2 and VSV-eGFP were generated and rescued as described previously (Case et al., 2020; Whelan et al., 1995). Briefly, BSRT7/5 cells (Buchholz et al., 1999) were infected with vaccinia virus encoding the bacteriophage T7 RNA polymerase (vTF7-3) (Fuerst et al., 1986) and subsequently transfected with plasmids encoding VSV N, P, L, G, and an antigenome copy of the viral genome under control of the T7 promoter. Rescue supernatants were collected 56 to 72 h post-transfection, clarified by centrifugation (5 min at 1,000 x g), and filtered through a 0.22 μm filter. Virus clones were plaque-purified on Vero CCL81 cells containing 25 μg/ml of cytosine arabinoside (Sigma-Aldrich) in the agarose overlay, and plaques were amplified on Vero CCL81 cells. All infections for generating stocks were performed at 37°C for 1 h and at 34°C thereafter. Viral supernatants were harvested upon extensive cytopathic effect and clarified of cell debris by centrifugation at 1,000 x g for 5 min. Aliquots were maintained at -80°C.

#### Mouse experiments

Animal studies were carried out in accordance with the recommendations in the Guide for the Care and Use of Laboratory Animals of the National Institutes of Health. The protocols were approved by the Institutional Animal Care and Use Committee at the Washington University School of Medicine (assurance number A3381–01). Virus inoculations were performed under anesthesia that was induced and maintained with ketamine hydrochloride and xylazine, and all efforts were made to minimize animal suffering.

At four weeks of age, female BALB/c mice (Jackson Laboratory, 000651) were immunized with 10^6^ PFU of VSV-eGFP-SARS-CoV-2 or VSV-eGFP via the intraperitoneal route. Where indicated, mice were boosted with homologous virus at 4 weeks post-priming. Three weeks post-priming or boosting mice were administered 2 mg of anti-Ifnar1 mAb (MAR1-5A3 (Sheehan et al., 2006), Leinco) via intraperitoneal injection. One day later, mice were administered 2.5 × 10^8^ PFU of mouse codon-optimized AdV-hACE2 (Hassan et al., 2020) via intranasal administration. Five days later, vaccinated mice were challenged with 3 × 10^5^ PFU of SARS-CoV-2 via intranasal administration. Passive transfer experiments were conducted as described above but using ten-week-old female BALB/c mice. Pooled immune sera were administered 24 h prior to SARS-CoV-2 challenge. For each immunization (prime or boost), serum from individual mice was collected twice (at days 14 and 22) and pooled.

### METHOD DETAILS

#### Gradient purification of recombinant viruses

To generate high titer stocks of VSV-eGFP and VSV-eGFP-SARS-CoV-2, viruses were grown on BSRT7/5 cells at an MOI of 3 or 1, respectively. To generate VSV-eGFP-SARS-CoV-2, BSRT7/5 cells were transfected with pCAGGS-VSV-G in Opt-MEM (Gibco) using Lipofectamine 2000 (Invitrogen) and subsequently infected 8 to 12 h later with VSV-eGFP-SARS-CoV-2 at an MOI of 0.01 in DMEM containing 2% FBS and 20 mM HEPES pH 7.7. This VSV G decorated VSV-eGFP-SARS-CoV-2 was titrated by plaque assay and used for a larger scale infection as described above. Cell supernatants were collected after 48 h and clarified by centrifugation at 1,000 x g for 7.5 min. Supernatants were concentrated using a Beckman Optima L-100 XP ultracentrifuge (22,800 RPM x 90 min in a 70Ti fixed-angle rotor). Pellets were resuspended in 100 mM NaCl, 10 mM Tris pH 7.4, 1 mM EDTA (NTE) at 4°C overnight, and virus banded on a 15-45% sucrose-NTE gradient (35,000 rpm x 3 h in a SW-41Ti swinging-bucket rotor). Virus was extracted by side puncture of tubes, recovered by ultracentrifugation (22,800 RPM x 90 min in a 70Ti fixed-angle rotor) and resuspended in NTE at 4°C overnight. The VSV-eGFP was purified similarly.

#### Measurement of viral burden

Mouse tissues were weighed and homogenized with sterile zirconia beads in a MagNA Lyser instrument (Roche Life Science) in 1 mL of DMEM media supplemented to contain 2% heat-inactivated FBS. Tissue homogenates were clarified by centrifugation at 10,000 rpm for 5 min and stored at -80°C. RNA was extracted using MagMax mirVana Total RNA isolation kit (Thermo Scientific) and a Kingfisher Flex extraction machine (Thermo Scientific). Infectious viral titers in lung homogenates were determined by plaque assays on Vero-furin cells. Viral RNA levels were determined by RT-qPCR as described (Hassan et al., 2020) and normalized to tissue weight.

#### Cytokine analysis

Total RNA was isolated from lung homogenates as described above and DNAase treated. cDNA was generated using the HighCapacity cDNA Reverse Transcription kit (Thermo Scientific) with the addition of RNase inhibitor according to the manufacturer’s instructions. Cytokine and chemokine expression were determined using TaqMan Fast Universal PCR master mix (Thermo Scientific) with commercially available primer/probe sets specific for *IFN-γ* (IDT: Mm.PT.58.41769240), *IL-6* (Mm.PT.58.10005566), *IL-1β* (Mm.PT.58.41616450), *TNF-α* (Mm.PT.58.12575861), *CXCL10* (Mm.PT.58.43575827), *CCL2* (Mm.PT.58.42151692), *CCL5* (Mm.PT.58.43548565), *CXCL11* (Mm.PT.58.10773148.g), *IFN-β* (Mm.PT.58.30132453.g), and *IFNλ-2/3* (Thermo Scientific Mm04204156_gH). All results were normalized to *GAPDH* (Mm.PT.39a.1) levels and the fold-change for each was determined using the 2^-Δ;Δ;Ct^ method comparing SARS-CoV-2 infected mice to naïve controls.

#### Histology and *in situ* hybridization

Mice were euthanized, and tissues were harvested prior to lung inflation and fixation. The right lung was inflated with approximately 1.2 mL of 10% neutral buffered formalin using a 3-mL syringe and catheter inserted into the trachea. To ensure fixation of virus, inflated lungs were kept in a 40-mL suspension of neutral buffered formalin for 7 days before further processing. Tissues were paraffin-embedded and sections were subsequently stained with hematoxylin and eosin. RNA *in situ* hybridization was performed using the RNAscope 2.5 HD Assay (Brown Kit) according to the manufacturer’s instructions (Advanced Cell Diagnostics). Briefly, sections were deparaffinized and treated with H_2_O_2_ and Protease Plus prior to RNA probe hybridization. Probes specifically targeting SARS-CoV-2 S sequence (cat no 848561) were hybridized followed by signal amplification and detection with 3,3′-Diaminobenzidine. Tissues were counterstained with Gill’s hematoxylin and an uninfected mouse was stained in parallel and used as a negative control. The lung pathology was evaluated, and representative photomicrographs were taken of stained slides under investigator-blinded conditions. Tissue sections were visualized using a Nikon Eclipse microscope equipped with an Olympus DP71 color camera or a Leica DM6B microscope equipped with a Leica DFC7000T camera.

#### Neutralization assay

Serial dilutions of mouse sera were incubated with 10^2^ focus-forming units (FFU) of SARS-CoV-2 for 1 h at 37°C. Antibody-virus complexes were added to Vero E6 cell monolayers in 96-well plates and incubated at 37°C for 1 h. Subsequently, cells were overlaid with 1% (w/v) methylcellulose in MEM supplemented with 2% FBS. Plates were harvested 30 h later by removing overlays and fixed with 4% PFA in PBS for 20 min at room temperature. Plates were washed and sequentially incubated with 1 mg/mL of CR3022 (PMID: 32245784) anti-S antibody and HRP-conjugated goat anti-human IgG in PBS supplemented with 0.1% saponin and 0.1% bovine serum albumin. SARS-CoV-2-infected cell foci were visualized using TrueBlue peroxidase substrate (KPL) and quantitated on an ImmunoSpot microanalyzer (Cellular Technologies). Data were processed using Prism software (GraphPad Prism 8.0).

#### ELISA

6-well Maxisorp plates were coated with 2 ug/mL of either SARS-CoV-2 Spike, RBD, NP, or ORF8 proteins in 50□mM Na_2_CO_3_ (70□μL) overnight at 4 °C. Plates were then washed with PBS + 0.05% Tween-20 and blocked with 200 µL 1X PBS + 0.05% Tween-20 + 1% BSA + 0.02% sodium azide for 2 h at room temperature (RT). Serum samples were serially diluted (1:3) starting at either 1:100 dilution (day 22 samples) or 1:30 dilution (day 8 post-infection samples) in blocking buffer. Diluted samples were then added to washed plates (50□μL/well) and incubated for 1 h at RT. Bound IgG was detected using HRP-conjugated goat anti-mouse IgG (at 1:2000) or bound IgM was detected using biotin-conjugated anti-mouse IgM (at 1:10000), followed by streptavidin-HRP (at 1:5000). Following a 1 h incubation, washed plates were developed with 50 µL of 1-Step Ultra TMB-ELISA, quenched with 2 M sulfuric acid, and the absorbance was read at 450□nm.

### QUANTIFICATION AND STATISTICAL ANALYSIS

Statistical significance was assigned when *P* values were < 0.05 using Prism Version 8 (GraphPad) and tests are indicated in the relevant Figure legends.

